# The putative causal effect of type 2 diabetes in risk of cataract: a Mendelian randomization study in East Asian

**DOI:** 10.1101/2021.02.08.430342

**Authors:** Haoyang Zhang, Xuehao Xiu, Angli Xue, Yuedong Yang, Yuanhao Yang, Huiying Zhao

## Abstract

**Background:** The epidemiological association between type 2 diabetes and cataract has been well-established. However, it remains unclear whether the two diseases share a genetic basis, and if so, whether this reflects a causal relationship.

**Methods:** We utilized East Asian population-based genome-wide association studies (GWAS) summary statistics of type 2 diabetes (*N*_case_=36,614, *N*_control_=155,150) and cataract (*N*_case_=24,622, *N*_control_=187,831) to comprehensively investigate the shared genetics between the two diseases. We performed 1. linkage disequilibrium score regression (LDSC) and heritability estimation from summary statistics (ρ-HESS) to estimate the genetic correlation and local genetic correlation between type 2 diabetes and cataract; 2. multiple Mendelian randomization (MR) analyses to infer the putative causality between type 2 diabetes and cataract; and 3. Summary-data-based Mendelian randomization (SMR) to identify candidate risk genes underling the causality.

**Results:** We observed a strong genetic correlation (*r*_g_=0.58; p-value=5.60×10^−6^) between type 2 diabetes and cataract. Both ρ-HESS and multiple MR methods consistently showed a putative causal effect of type 2 diabetes on cataract, with estimated liability-scale MR odds ratios (ORs) at around 1.10 (95% confidence interval [CI] ranging from 1.06 to 1.17). In contrast, no evidence supports a causal effect of cataract on type 2 diabetes. SMR analysis identified two novel genes *MIR4453HG* (*β*_SMR_=−0.34, p-value=6.41×10^−8^) and *KCNK17* (*β*_SMR_=−0.07, p-value=2.49×10^−10^), whose expression levels were likely involved in the putative causality of type 2 diabetes on cataract.

**Conclusions:** Our results provided robust evidence supporting a causal effect of type 2 diabetes on the risk of cataract in East Asians, and posed new paths on guiding prevention and early-stage diagnosis of cataract in type 2 diabetes patients.

**Key Messages:**

- We utilized genome-wide association studies of type 2 diabetes and cataract in a large Japanese population-based cohort and find a strong genetic overlap underlying the two diseases.
- We performed multiple Mendelian randomization models and consistently disclosed a putative causal effect of type 2 diabetes on the development of cataract.
- We revealed two candidate genes *MIR4453HG* and *KCNK17* whose expression levelss are likely relevant to the causality between type 2 diabetes and cataract.
- Our study provided theoretical fundament at the genetic level for improving early diagnosis, prevention and treatment of cataract in type 2 diabetes patients in clinical practice

## Introduction

Type 2 diabetes is one of the most prevalent chronic diseases in East Asians^1^, and cataract is a major cause of vision impairment among patients with type 2 diabetes^2^. Previous studies^2, 3^ have revealed a strong phenotypic association between type 2 diabetes and cataract for East Asians. For instance, Foster et al. conducted a cross-sectional study of 1,206 Singapore Chinese and found patients with diabetes had higher risks for obtaining cortical cataract^3^. Another Asian population-based study recruited 10,033 participants and identified diabetes as a significant risk factor for elevating incidence of cataract surgery^2^.

The phenotypic association between type 2 diabetes and cataract could be partially explained by their shared genetics^4, 5^. As pieces of evidence for disclosing their shared genetics, Lee et al.^4^ analyzed a Hong Kong Chinese cohort and found cataract is common in patients with type 2 diabetes who carried microsatellite polymorphism around aldose reductase-related genes. Lin et al.^5^ identified multiple candidate genes that had significantly different expression levels in the type 2 diabetes patients with higher Lens Opacities Classification System (LOCS) score (i.e., a system used to grade age-related cataract^6^), comparing to the patients with zero or minor LOCS score. However, the magnitude of the genetic association between type 2 diabetes and cataract remains unclear, as does the problem of whether their genetic association reflects a causal relationship.

Traditional methods estimate the shared genetics by comparing the concordance between monozygotic and dizygotic twins^7^, and establish causal conclusions using the randomized controlled trials (RCTs)^8^, a widely accepted gold standard for causal inference. However, these methods are occasionally limited or impracticable due to their own methodological weakness, such as the laborious data collection process and unethical study design. With the development of genome-wide association studies (GWAS) during past decades, some alternatively feasible statistical methods have been proposed to estimate the shared genetics between focal traits directly using the GWAS summary data^9^. For instance, Bulik-Sullivan et al.^10^ developed a technique named linkage disequilibrium score regression (LDSC) to estimate the contributions of polygenic genetic effects for a focal trait (i.e., single-trait heritability) and the magnitude of shared genetic overlap underlying two traits (i.e., cross-trait genetic correlation). Shi et al.^11^ extended LDSC and proposed heritability estimation from summary statistics (ρ-HESS), a method to quantify the local single-trait heritability and cross-trait genetic correlation from approximately LD-independent genomic regions. For pair of traits with significant genetic correlation, Mendelian randomization (MR)^12^ methods are capable of inferring the potential genetic causal relationship between traits using single nucleotide polymorphisms (SNPs) as instruments. To further investigate any putative functional genes underlying the susceptibility to a trait, Zhu et al.^13^ proposed summary data-based Mendelian randomization (SMR), which is an approach to identify gene expressions in an association with a target trait, by integrating GWAS summary data with expression quantitative trait loci (eQTL) summary data.

In this study, we leveraged the large East Asian population-based GWAS summary statistics of type 2 diabetes and cataract from BioBank Japan Project (BBJ)^14^ to comprehensively investigate the shared genetics between the two diseases. We applied LDSC, ρ-HESS, and seven MR or MR-equivalent approaches to estimate the genetic correlation, local genetic correlation, and potential genetic causality between type 2 diabetes and cataract, respectively. We also conducted SMR to the single-trait GWAS (i.e., type 2 diabetes, cataract) and cross-trait GWAS meta-analyses of type 2 diabetes and cataract to explore candidate genes involved in the causality between two diseases. A brief overview of our study is summarized in Fig. 1.

**Figure 1.**
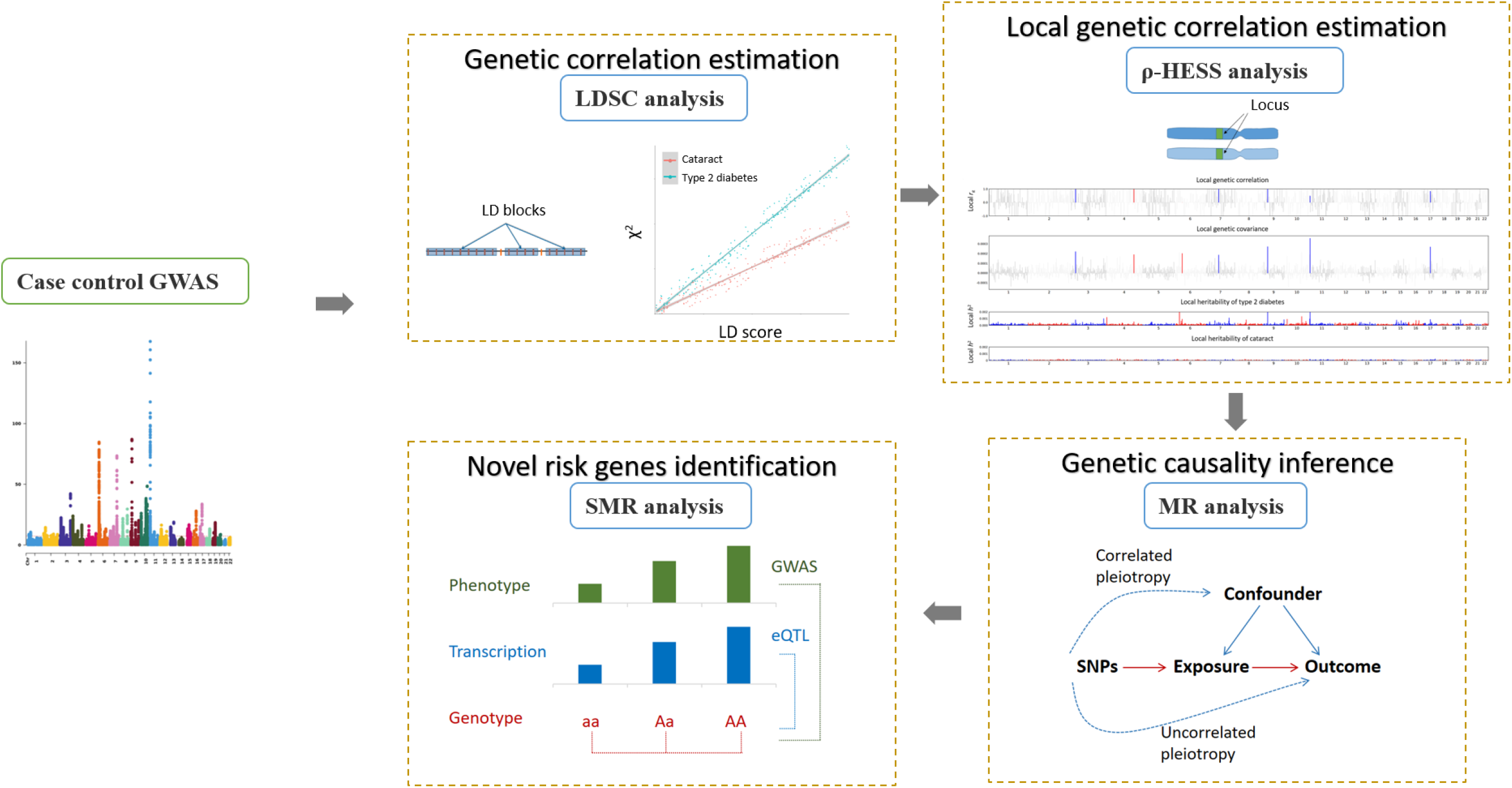
Outline of statistical analysis performed in our study, including four sections: 1. top left, measuring the genetic correlation between type 2 diabetes and cataract using LDSC; 2. top right, measuring the local genetic correlation between type 2 diabetes and cataract using ρ-HESS; 3. bottom right, investigating the potential causal relationship between type 2 diabetes and cataract using seven MR or MR-equivalent approaches. 4. bottom left, identifying candidate risk genes underlying the causality between type 2 diabetes and cataract. LDSC: linkage disequilibrium score regression; ρ-HESS: Heritability Estimation from Summary Statistics; MR: Mendelian randomization. SMR: summary data-based Mendelian randomization.

## Research Design and Methods

### GWAS Data Source

We downloaded the GWAS summary statistics of type 2 diabetes^15^ and cataract^16^ from the BBJ (http://jenger.riken.jp/en/), a database with common diseases, complex traits, and demographic and genotype data from ~200,000 Japanese individuals (53.10% male; average baseline age at 62.70 for men and 61.50 for women^14^). Both type 2 diabetes and cataract were diagnosed by physicians. The GWAS of type 2 diabetes was generated using a fixed-effect inverse-variance meta-analysis via METAL^17^, comprising 36,614 cases and 155,150 controls of four Japanese ancestry-based cohorts^15^. The GWAS of cataract was generated from 24,622 cases and 187,831 controls using a linear mixed model via SAIGE^18^, adjusted by age, sex, and top five principal components^16^. Both GWAS were based on the hg19 coordinate.

### LDSC of single-trait heritability and cross-trait genetic correlation

We applied LDSC^10, 19^ to estimate the liability-scale heritability (*h*^2^) of type 2 diabetes and cataract as well as their genetic correlation (*r*_g_). GWAS summary statistics were filtered according to the HapMap3 reference^20^. SNPs were excluded if they were strand-ambiguous, had minor allele frequency <0.01, or located within the major histocompatibility complex (MHC) region (chromosome 6: 28,477,797–33,448,354) due to the complicated LD structure in this region^21^. The LD scores were pre-computed based on the 481 East Asians in 1000 Genomes (https://alkesgroup.broadinstitute.org/LDSCORE/). Univariate LDSC was performed to estimate the liability-scale *h^2^* of type 2 diabetes and cataract, assuming the population and sample prevalence at 7.50%^15^ and 19.10%^15^ for type 2 diabetes, and 0.09%^22^ and 11.59%^16^ for cataract, respectively. Bivariate LDSC was utilized to estimate the genetic correlation (i.e., *r*_g_) between type 2 diabetes and cataract with and without a constrained intercept, which is designed to reduce the bias from population stratification. A significant *r*_g_ was determined with p-value <0.05.

### ρ-HESS of local genetic correlation

To explore whether type 2 diabetes had significant genetic overlap with cataract in some specific independent genomic regions, we performed ρ-HESS^11^ to estimate the local genetic correlations between type 2 diabetes and cataract according to the hg19-based 1000 Genomes East Asian reference. A total of 1,439 approximately LD-independent genomic regions (with the exclusion of the MHC region)^23^ were utilized in our analysis. The regions were excluded if the estimated local single-trait heritability was negative because of the insufficient study power. The estimated local genetic correlations were divided into four regional types: 1. the regions harboring significant type 2 diabetes-specific SNPs (i.e., ‘type 2 diabetes-specific’); 2. the regions harboring significant cataract-specific SNPs (i.e., ‘cataract-specific’); 3. the regions harboring shared SNPs significantly associated with both type 2 diabetes and cataract (i.e., ‘intersection’); and 4. other regions (i.e., ‘neither’). Three GWAS p-value thresholds, 5×10^−8^, 1×10^−5^, and 1×10^−3^, were used to define the significant SNPs. For these four regional types occupied by more than 10 regions, we calculated the mean and standard error of local genetic correlations within each type. A potential causal effect of type 2 diabetes on cataract is suggested if the average local genetic correlation at type 2 diabetes-specific regions and cataract-specific regions were significantly and non-significantly different from zero, respectively. The opposite is true for inferring potential causal effect of cataract on type 2 diabetes. Besides, the existence of pleiotropic effect may be implicated if there is a non-zero average local genetic correlation at intersection regions.

### MR analyses for genetic causality inference

The causal relationship between type 2 diabetes and cataract was evaluated by six MR approaches (i.e., inverse variance weighted [IVW] model^24^, MR-Egger model^25^, generalized summary-data-based Mendelian randomization [GSMR]^26^, weighed median model^27^, and weighted mode model^28^, and the causal analysis using summary effect estimates [CAUSE]^29^) and one MR-equivalent latent causal variable (LCV) model^30^. Multiple methods were employed because they have different assumptions on horizontal pleiotropy, a term defined as the instrumental SNPs with effects on both exposure and outcome through non-causal pathways^12^. Horizontal pleiotropy is a potential confounding factor for inferring causality and can be divided into uncorrelated pleiotropy if the instrumental SNPs influence exposure and outcome via independent mechanisms, and correlated pleiotropy if the instrumental SNPs affect exposure and outcome through shared factors^12^. The consistent results of multiple MR methods are expected to effectively minimize the impact of horizontal pleiotropy^31^ from putative causality and thus reduce the false-positive rate.

Among the seven methods, IVW measures the causal effect by integrating ratios of variant effects (ratio estimates) between exposure and outcome, assuming a balanced uncorrelated pleiotropy (with mean zero) and no correlated pleiotropy. MR-Egger assumes no correlated pleiotropy and a non-zero uncorrelated pleiotropy, which adds an extra intercept compared to IVW to represent the magnitude of uncorrelated pleiotropy. GSMR assumes the presence of uncorrelated pleiotropy and excludes such effect by outlier removal using heterogeneity in dependent instrument (HEIDI) approach. The weighted median model assumes the proportion of pleiotropic (both uncorrelated and correlated) instrumental SNPs is less than half, and calculates the causal effect using the weighted median of the SNP ratio. The weighted mode model greatly loosens the assumptions on uncorrelated and correlated pleiotropy and measures the causal effect only from the most frequent (the mode) SNP set with consistent effect. At least 10 independent instrumental SNPs are required for maintaining study power using these five MR methods. Independent instrumental SNPs are selected from the exposure-specific genome-wide significant (GWAS p-value<5×10^−8^) SNPs that are also merged with outcome GWAS, and then clumped by LD *r*^2^<0.05 within 1,000 kb window using PLINK version 1.9^32^ according to the reference genome of 1000 Genomes East Asian^33^. If the number of independent instrumental SNPs was less than 10, we selected the ‘proxy’ instrumental SNPs by relaxing the exposure GWAS p-value threshold to 1×10^−5^ to maintain study power. The LCV model^30^ is a MR-equivalent method that assumes the genetic correlation between two traits is mediated by a latent variable, and distinguishes causality from uncorrelated and correlated pleiotropy by measuring the genetic causality proportion (GCP) using all genetic variants^30^. CAUSE^29^ is a method that is more powerful and sensitive to identify the causality from uncorrelated and correlated pleiotropy compared to other models. CAUSE increases MR detection power by recruiting more approximately independent instrumental SNPs with GWAS p-value <1×10^−3^ and pruned by LD *r^2^* <0.1. CAUSE also provides an expected log pointwise posterior density (ELPD) test to compare the overall fitness among a causal model (i.e., instrumental SNPs act on exposure and outcome through a causal pathway and shared factors), a sharing model (i.e., instrumental SNPs act on exposure and outcome only through shared factors), and a null model (i.e., no causal pathway or shared factors underlying exposure and outcome).

We performed these models using R packages “*cause*” (version: 1.0.0), “*LCV*”, “*gsmr*” (version: 1.0.9) and “*TwoSampleMR*” (version: 0.5.4). Any instrumental SNPs located within the MHC region were excluded^21^. A significant causal relationship was determined if the causal effect estimates were consistent and significant at the Bonferroni-corrected level (with p-value<0.05/13≈3.85×10^−3^, including six bi-directional MR methods and LCV model). The causal effects (i.e., *β*) were converted from logit-scale to liability-scale using the method described by Byrne et al^34^:

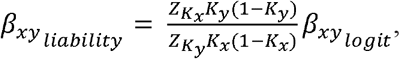

where *K*_*x*_ and *K*_*y*_ are the population prevalence of exposure and outcome, and 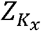 and 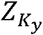 are the height of standard normal distribution at such prevalence. We assumed the population prevalence for type 2 diabetes and cataract are 7.5%^15^ and 0.09%^22^ that are same as the population prevalence used in estimating the liability heritability. The liability *β* was then transformed into odds ratio (OR).

### SMR analysis to identify candidate genes underling the genetic causality

When a causal relationship between type 2 diabetes and cataract is established, we applied SMR^13^ to identify candidate risk genes underling the causality between the two diseases. SMR leverages GWAS and eQTL summary statistics to explore the association between gene expression and a target disease or trait using a MR equivalent analysis. SMR further utilizes HEIDI-outlier test to check if the significant association between gene expression and the target trait/disease is due to the causality (i.e., causal SNPs drive disease by regulating gene expression levels) or pleiotropy (i.e., causal SNPs influences both disease and gene expression via shared effects) rather than linkage (i.e., different causal SNPs in LD influences disease and gene expression, respectively).

In our study, we performed SMR using cis-eQTLgen summary data (19,250 expression probes in blood; *URL*: https://eqtlgen.org/cis-eqtls.html)^35^ to the single-trait GWAS (i.e., type 2 diabetes, cataract) and cross-trait GWAS of type 2 diabetes and cataract generated by inverse-variance-based meta-analysis via METAL^36^. Significant gene expressions due to causality or pleiotropy were determined if with a study-wise Bonferroni-corrected SMR p-value <0.05/19,250/3 ≈ 8.66×10^−7^ and a HEIDI-outlier p-value >0.05 calculated from minimum 10 SNPs. To identify any candidate risk genes possibly involved in the causality between type 2 diabetes and cataract, we focused on the gene expressions that were significantly associated with cross-trait meta-analysis of type 2 diabetes and cataract, but not with the original single-trait disease (i.e., type 2 diabetes or cataract).

## Results

### Strong genetic association between type 2 diabetes and cataract

As shown in Table 1, the estimated SNP-based liability-scale *h^2^* for type 2 diabetes and cataract were 21.47% (standard error [SE]=1.17%, p-value=3.28×10^−75^) and 1.63% (SE=0.13%, p-value=4.60×10^−36^) with constrained LDSC intercept, respectively. These *h^2^* decreased to 15.12% (SE=1.37%, p-value=2.55×10^−28^) and 0.54% (SE=0.19%, p-value=4.48×10^−3^) without constrained LDSC intercept, suggesting the mild inflations in both diseases GWAS. We then performed the bivariate LDSC with and without constrained intercept, and identified the significant genetic correlation *r*_g_ between type 2 diabetes and cataract at 0.28 (SE=0.05, p-value=3.25×10^−9^) and 0.58 (SE=0.13, p-value=5.60×10^−6^), respectively, indicating strong shared genetics between type 2 diabetes and cataract.

**Table 1.** Heritability of type 2 diabetes and cataract and their genetic correlation estimated by LDSC.

|  |  | Constrained intercept |  | Unconstrained intercept |  |
| --- | --- | --- | --- | --- | --- |
|  |  | Type 2 diabetes | Cataract | Type 2 diabetes | Cataract |
| Single-trait | Heritability ($h^2 \pm \text{SE}$) | 0.215±0.012 | 0.016±0.001 | 0.151±0.014 | 0.005±0.002 |
| LDSC | $P_{h2}$ | 3.28$\times 10^{-75}$ | 4.60$\times 10^{-36}$ | 2.55$\times 10^{-28}$ | 4.48$\times 10^{-3}$ |
| Cross-trait | Genetic correlation ($r_g \pm \text{SE}$) | 0.284±0.048 | | 0.575±0.127 | |
| LDSC | $P_{rg}$ | 3.25$\times 10^{-9}$ | | 5.60$\times 10^{-6}$ | |

### ρ-HESS analyses of local genetic correlations

We conducted ρ-HESS to estimate the local heritability of type 2 diabetes and cataract (detailed results in Table S1 and Fig. 2A). We also estimated the local genetic covariance and correlation between type 2 diabetes and cataract in 824 regions (detailed in Table S1) after excluding the regions with negative local heritability. As shown in Table 2, we identified six genomic regions at a nominal significance level (p-value<0.05) from different chromosomes, with estimated local *r*_g_ at [0.48, 1).

**Table 2.** Local heritability of type 2 diabetes and cataract and their genetic covariance/correlation estimated by ρ**-HESS**.

| Region |  |  |  | Type 2 diabetes |  |  | Cataract |  |  | Cross-trait |  |  |  |
| --- | --- | --- | --- | --- | --- | --- | --- | --- | --- | --- | --- | --- | --- |
| Chr | Start | End | $N_{\text{SNP}}$ | local $h^2$ | $\text{SE}_{h^2}$ | $P_{h^2}$ | local $h^2$ | $\text{SE}_{h^2}$ | $P_{h^2}$ | local covariance | $\text{SE}_{\text{covariance}}$ | local $r_g^*$ | $P_{rg}$ |
| 3 | 22647520 | 25460047 | 5859 | $8.19 \times 10^{-4}$ | $1.74 \times 10^{-4}$ | $1.24 \times 10^{-6}$ | $3.25 \times 10^{-5}$ | $1.43 \times 10^{-4}$ | $4.10 \times 10^{-1}$ | $2.32 \times 10^{-4}$ | $1.05 \times 10^{-4}$ | 1 | $2.77 \times 10^{-2}$ |
| 4 | 152782184 | 154122370 | 1547 | $5.339 \times 10^{-4}$ | $1.60 \times 10^{-4}$ | $4.23 \times 10^{-4}$ | $7.23 \times 10^{-5}$ | $1.46 \times 10^{-4}$ | $3.10 \times 10^{-1}$ | $2.00 \times 10^{-4}$ | $9.97 \times 10^{-5}$ | 1 | $4.49 \times 10^{-2}$ |
| 7 | 68458764 | 69083424 | 1102 | $4.58 \times 10^{-4}$ | $1.51 \times 10^{-4}$ | $1.22 \times 10^{-3}$ | $2.34 \times 10^{-5}$ | $1.37 \times 10^{-4}$ | $4.32 \times 10^{-1}$ | $1.96 \times 10^{-4}$ | $9.38 \times 10^{-5}$ | 1 | $3.70 \times 10^{-2}$ |
| 9 | 21938460 | 23424761 | 2356 | $2.01 \times 10^{-3}$ | $2.24 \times 10^{-4}$ | $1.62 \times 10^{-19}$ | $4.17 \times 10^{-5}$ | $1.44 \times 10^{-4}$ | $3.86 \times 10^{-1}$ | $2.86 \times 10^{-4}$ | $1.29 \times 10^{-4}$ | 0.98 | $2.64 \times 10^{-2}$ |
| 11 | 2739889 | 4466102 | 2594 | $3.24 \times 10^{-3}$ | $2.66 \times 10^{-4}$ | $1.74 \times 10^{-34}$ | $1.88 \times 10^{-4}$ | $1.53 \times 10^{-4}$ | $1.09 \times 10^{-1}$ | $3.76 \times 10^{-4}$ | $1.51 \times 10^{-4}$ | 0.48 | $1.30 \times 10^{-2}$ |
| 17 | 34395061 | 36495389 | 1844 | $8.83 \times 10^{-4}$ | $1.77 \times 10^{-4}$ | $3.04 \times 10^{-7}$ | $1.30 \times 10^{-4}$ | $1.50 \times 10^{-4}$ | $1.92 \times 10^{-1}$ | $2.83 \times 10^{-4}$ | $1.09 \times 10^{-4}$ | 0.83 | $9.26 \times 10^{-3}$ |

\*The estimates of local genetic correlation may be less than -1 or greater than 1 due to the insufficient study power. ρ-HESS caps these estimates at -1 or 1.

**Figure 2.**
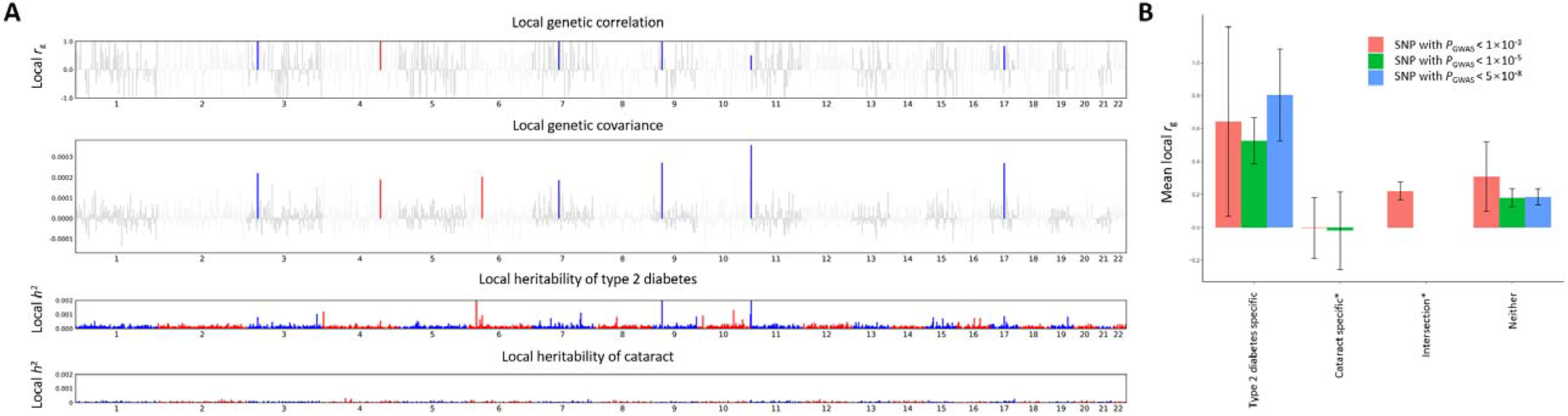
A. The local heritability of type 2 diabetes and cataract as well as their genetic covariances and genetic correlations from approximately LD-independent genomic regions estimated by ρ-HESS. For plots of ‘local heritability’, the blue or red bars represent the genomic regions on odds or even chromosome. For the plots of ‘local genetic correlation’ and ‘local genetic covariance’, the blue or red bars represent the genomic regions on odds or even chromosome showing nominal significant (p-value <0.05) local genetic correlation/covariance between type 2 diabetes and cataract. B. The average local genetic correlation between type 2 diabetes and cataract in four regional types (i.e., ‘type 2 diabetes-specific’, ‘cataract-specific’, ‘intersection’, and ‘neither’) harboring risk SNPs with GWAS p-value <1×10^−3^ (colored in red; *N*_region_=519, 87, 685, and 146), <1×10^−5^ (colored in green; *N*_region_=232, 29, 5, and 1171) and <5×10^−8^ (colored in blue; *N*_region_=84, 1, 0, and 1352), respectively. Error bars represent the 95% confidence intervals (CIs) of the estimates. ρ-HESS: Heritability Estimation from Summary Statistics.

We further investigated the distribution of local genetic correlations in four regional types. As shown in Fig. 2B, regions harboring type 2 diabetes-specific SNPs were identified with average local *r*_g_ significantly higher than zero. In reverse, regions harboring cataract-specific SNPs showed a non-significant average local *r*_g_ close to zero. Therefore, the distribution of local *r*_g_ revealed by ρ-HESS suggested a potential putative causal relationship of type 2 diabetes on cataract. Besides, the average local *r*_g_ from the ‘intersection’ regions harboring shared significant SNPs with GWAS p-value<1×10^−3^ was estimated at 0.22 (SE=0.03, p-value=1.19×10^−15^), suggesting the mild pleiotropic effects of ‘less significant’ genetic variants may be underlying type 2 diabetes and cataract, while we cannot further distinguish the causality of type 2 diabetes on cataract from such pleiotropy here using ρ-HESS.

### Putative causality of type 2 diabetes on cataract

Application of seven MR or MR-equivalent methods consistently detected a causal effect of type 2 diabetes on cataract at Bonferroni-corrected significance level (p-value<3.85×10^−3^), detailed in Table S2. In reverse, there was no or modest evidence for a causal effect of cataract on type 2 diabetes. As shown in Fig. 3, six MR methods provided consistent evidence for a causal effect of type 2 diabetes on cataract with estimated liability-scale ORs ranging from 1.07 to 1.13, under the assumption of the population prevalence at 7.50% of type 2 diabetes and 0.09% of cataract, respectively. These results indicated that individuals with type 2 diabetes had approximately 1.06 to 1.17 times of risks for developing cataract compared to healthy individuals. Besides, LCV provided a GCP at 0.87 (p-value=1.54×10^−9^), suggesting that the strong genetic correlation between type 2 diabetes and cataract can be largely explained by the causality of type 2 diabetes on cataract. Remarkably, our putative causality of type 2 diabetes on cataract was less likely to be influenced by the horizontal pleiotropy because of the close-to-zero MR-Egger intercept (−0.002, p-value =0.47) and the better model fitness of causal model (with non-significant effects of correlated pleiotropy [*η*=−0.02, 95% CI=−0.53 to 0.30] and uncorrelated pleiotropy [*q*=0.04, 95% CI=0 to 0.25]) compared to the sharing model (ELPD p-value=8.80×10^−3^) and the null model (ELPD p-value=4.69×10^−13^) revealed by CAUSE (Fig. S1).

**Figure 3.**
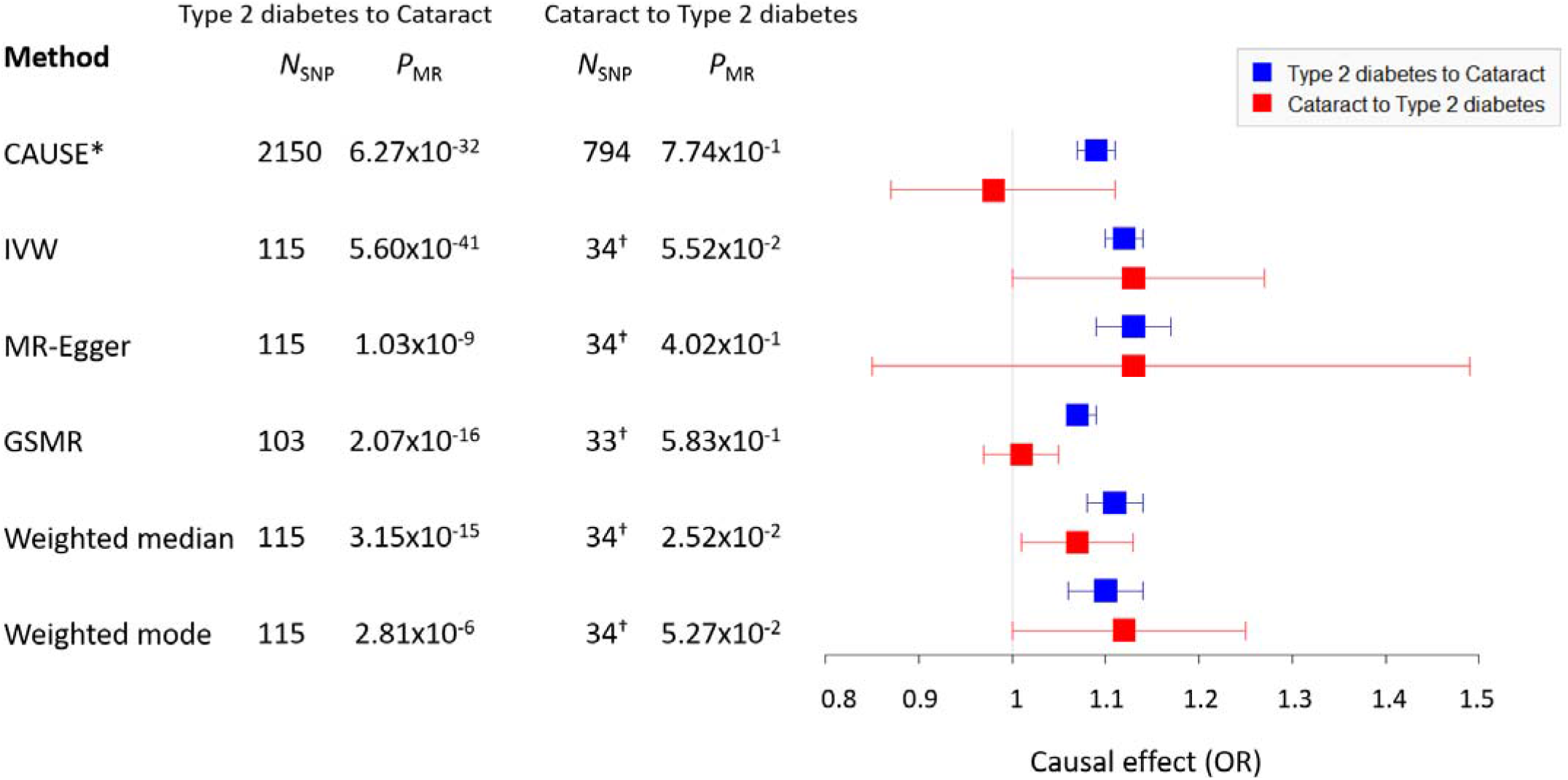
The bi-directional causal effect (in liability-scale odds ratio [OR]) between type 2 diabetes and cataract estimated by six MR methods. Error bars represent the 95% confidence intervals (CIs) of the estimates. Blue: type 2 diabetes as exposure and cataract as outcome, i.e., ‘Type 2 diabetes to Cataract’; Red: cataract as exposure and type 2 diabetes as outcome, i.e., ‘Cataract to Type 2 diabetes’. CAUSE: causal analysis using summary effect estimates; GSMR: generalized summary-data-based Mendelian randomization; IVW: inverse variance weighted. ^*^CAUSE recruited independent instrumental SNPs with GWAS p-value <1×10^−3^. ^†^IVW, MR-Egger, GSMR, weighted median, and weighted mode recruited ‘proxy’ independent instrumental SNPs with GWAS p-value <1×10^−5^.

### Two candidate genes likely involved in the causality of type 2 diabetes on cataract

As shown in Table S3, we performed a cross-trait meta-analysis of type 2 diabetes and cataract using METAL, and identified 9 independent ‘novel’ SNPs that were associated with cross-trait of type 2 diabetes and cataract but not with the original GWAS of type 2 diabetes or cataract. Next, we applied SMR to the single-trait GWAS and the cross-trait meta-analysis GWAS of type 2 diabetes and cataract, and identified two candidate risk genes (Table 3), *MIR4453HG* (*β*_SMR_=−0.34; SMR p-value=6.41×10^−8^; HEIDI p-value=0.08 from 13 SNPs) and *KCNK17* (*β*_SMR_=−0.07; SMR p-value=2.49×10^−10^; HEIDI p-value=0.08 from 17 SNPs), whose expression levels were negatively associated (i.e., lower gene expression level increases the disease risk) with the susceptibility to co-morbid type 2 diabetes and cataract but not with the single-traits. These genes likely play crucial roles in the casual effects of type 2 diabetes on cataract.

**Table 3.** SMR identified two candidate genes involving in the causality between type 2 diabetes and cataract.

| Gene | Cross-trait Meta |  | Type 2 diabetes |  | Cataract |  |
| --- | --- | --- | --- | --- | --- | --- |
|  | *MIR4453HG* | *KCNK17* | *MIR4453HG* | *KCNK17* | *MIR4453HG* | *KCNK17* |
| Chr | 4 | 6 | 4 | 6 | 4 | 6 |
| Location of probe | 153,458,914 | 39,274,553 | 153,458,914 | 39,274,553 | 153,458,914 | 39,274,553 |
| Top SNP | rs4696318 | rs11753141 | rs10023327 | rs3734618 | rs10023327 | rs3734618 |
| Effect allele | A | A | T | G | T | G |
| Non-effect allele | G | G | A | A | A | A |
| $\beta_{\text{GWAS}} (\pm\text{SE})$ | -0.05±0.01 | 0.04±0.01 | -0.06±0.01 | 0.06±0.01 | -0.03±0.01 | 0.03±0.01 |
| $P_{\text{GWAS}}$ | 2.79$\times 10^{-14}$ | 1.84$\times 10^{-10}$ | 3.57$\times 10^{-12}$ | 9.56$\times 10^{-10}$ | 1.46$\times 10^{-2}$ | 5.18$\times 10^{-3}$ |
| $\beta_{\text{eQTL}} (\pm\text{SE})$ | 0.15±0.02 | -0.65±0.01 | 0.14±0.02 | -0.65±0.01 | 0.14±0.02 | -0.65±0.01 |
| $P_{\text{eQTL}}$ | 1.23$\times 10^{-14}$ | 0 | 1.10$\times 10^{-13}$ | 0 | 1.10$\times 10^{-13}$ | 0 |
| $\beta_{\text{SMR}} (\pm\text{SE})$ | -0.34±0.06 | -0.07±0.01 | -0.46±0.09 | -0.09±0.01 | -0.18±0.08 | -0.05±0.02 |
| $P_{\text{SMR}}$ | 6.41$\times 10^{-8}$ | 2.49$\times 10^{-10}$ | 3.82$\times 10^{-7}$ | 1.10$\times 10^{-9}$ | 2.03$\times 10^{-2}$ | 5.21$\times 10^{-3}$ |
| $P_{\text{HEIDI}}$ | $8.28 \times 10^{-2}$ | $7.90 \times 10^{-2}$ | $1.16 \times 10^{-1}$ | $1.05 \times 10^{-2}$ | $1.44 \times 10^{-1}$ | $8.81 \times 10^{-1}$ |
| $N_{\text{HEIDI\_SNP}}$ | 13 | 17 | 8 | 20 | 7 | 20 |

## Discussion

To our knowledge, this is the first study to quantify the genetic correlation and explore the potential causality between type 2 diabetes and cataract specifically using East Asian population-based GWAS summary statistics. Our results have highly enriched our current knowledge on the shared genetic architecture between type 2 diabetes and cataract.

Previously, researchers preferred to define co-occurrence of cataract and diabetes as a single outcome (i.e., diabetic cataract) and explored its genetics straightforwardly. For example, Lin et al.^5^ performed a GWAS using 758 Chinese cases with type 2 diabetic cataract and 649 healthy controls and identified 15 independent genome-wide significant SNPs, which are associated with blood sugar regulation and cataract development. Another study^37^ recruited 2,501 Scottish cases and 3,032 controls and found a significant role of rs2283290 in triggering diabetic cataract. Instead of using a single GWAS dataset with a small number of diabetic cataract patients, we leveraged large population-based GWAS summary statistics of type 2 diabetes and cataract, which is more powerful and provided robust evidence supporting the shared genetics between type 2 diabetes and cataract^3,4,38^.

Using ρ-HESS, we identified six genomic regions with a significant local genetic correlation between type 2 diabetes and cataract. Assuming these regions might contribute to the causal effect of type 2 diabetes on cataract, any SNPs or genes that are located within such regions and associated with type 2 diabetes and/or cataract risks are of great interest to understand the mechanisms underlying the regions. Therefore, we collected information from a total of 254 SNPs in ClinVar^39^ and genes in Malacards (supported by trustworthy sources or Cochrane based reviews^40^) for further analyses (Table S5). We identified gene *HNF1B* (hepatocyte nuclear factor 1*β*; Chromosome: 17: 36,046,434–36,105,096) and two SNPs (i.e., rs121918673 [Chromosome: 17: 36,061,127] and rs1555818071 [Chromosome: 17: 36,047,338]; located within gene *HNF1B*) that located within the significant genomic region on chromosome 17: 34,395,061–36,495,389 and reported to be associated with type 2 diabetes^41, 42^. In contrast, no SNPs or genes located in the ρ-HESS estimated significant genomic regions were found to be associated with cataract risk. Additionally, this result revealed a large proportion of shared genetics between type 2 diabetes and cataract were from the ‘type 2 diabetes-specific’ regions. These findings provided further evidence that the strong genetic correlation between type 2 diabetes and cataract is due to the type 2 diabetes-specific variants.

Application of seven MR and MR-equivalent methods provided consistent results for a causal effect of type 2 diabetes on cataract. Our findings raise an important clinical concern in prevention and early-diagnosis of cataract in patients with type 2 diabetes. We provided theoretical basis at genetic level for suggesting that assessing the development and severity of type 2 diabetes is likely yielding new targets for early-diagnosis of cataract, while further studies are required to pinpoint the potential aetiology underlying type 2 diabetes and cataract.

We also tried to replicate our findings in the European cohort using the European population-based publicly available GWAS summary statistics of type 2 diabetes^43^ (*N*_case_=62,892, *N*_control_=596,424) and cataract (*N*_case_=5,045, *N*_control_=356,096; UKB field ID: 20002; accessed from *URL*: http://www.nealelab.is/uk-biobank). However, LDSC analysis indicated a non-significant genetic correlation between the two diseases according to either European or East Asian reference (see Table S6). This result suggests the shared genetic variance between type 2 diabetes and cataract in East Asians may have strong genetic heterogeneity compared to Europeans. Future investigations are required for a better understanding of such difference.

To identify any blood-based biomarkers that may contribute to the causal effect of type 2 diabetes, we performed the multi-trait-based conditional & joint analysis (mtCOJO)^44^ to adjust both type 2 diabetes and cataract GWAS on each blood-based biomarker and then conducted post-mtCOJO MR analysis on adjusted type 2 diabetes and cataract GWAS (see Supplementary note, Table S7-9, and Fig. S2-3). We found that HbA1c (i.e., Hemoglobin A1c) may be involved in the causality of type 2 diabetes on cataract, standing in line with previous RCTs showing the impact of glycemic control on the prevention of ocular complications^45–47^. However, this result was possibly caused by the high genetic correlation between HbA1c and type 2 diabetes (*r*_g_=0.57 and 0.84 with and without constrained intercept) which may greatly decrease the heritability of type 2 diabetes and thus reduced the genetic correlation and putative causal relationship between type 2 diabetes and cataract. Future investigations should focus on this finding with the recruitment of a larger sample size.

We identified two candidate functional genes *MIR4453HG* and *KCNK17* that are likely relevant to the genetic causality between type 2 diabetes and cataract. Interestingly, both genes described a significant association with single-trait type 2 diabetes due to linkage (i.e., not passed HEIDI-outlier test), and then showed a more significant association with cross-trait type 2 diabetes and cataract due to causality or pleiotropy, further suggesting that cataract is likely an outcome triggered by the genetic mutations of type 2 diabetes. *MIR4453HG* is an IncRNA gene and located nearby some risk genes that have been reported to be associated with blood protein level (gene *ARFIP1*^48^) and lipoprotein cholesterol levels (gene *TRIM2*^49^). Both traits are highly relevant to the risk for type 2 diabetes^50,51^ and cataract^52,53^. *KCNK17* encoded a protein in the family of potassium channel^54^. The mutation of *KCNK17* may cause the abnormal opening of potassium channels and is associated with cardiovascular diseases (e.g., ischemic stroke and cerebral hemorrhage)^54^, which are known to be involved in the susceptibility to both type 2 diabetes^55^ and cataract^56^. These results provided novel insights on the genetic mechanisms underlying the causality between type 2 diabetes and cataract. Further wet-lab experiments were required to approve the roles of these two genes in increasing cataract risks in type 2 diabetes patients.

Our study has several limitations. First, the heritability of cataract was tiny with an estimate less than 2%, which may bias the estimate of genetic correlation between type 2 diabetes and cataract. Nevertheless, this effect should be negligible as the heritability of cataract is significantly different from zero. Secondly, the number of instrumental SNPs using cataract as exposure is less than 10. Instead, we selected the ‘proxy’ instrumental SNPs with p-value <1×10^−5^, which may violate assumptions of some MR methods (e.g., GSMR). However, the MR effects of these MR methods are highly consistent with CAUSE, suggesting the feasible application of using the ‘proxy’ instrumental SNPs. Thirdly, due to the limitation of our statistical models, we did not investigate the genetic contributions of the MHC region on the susceptibility to co-morbid type 2 diabetes and cataract, which possibly underestimated the shared genetic between the two diseases.

In summary, we provide robust evidence for a strong genetic association between type 2 diabetes and cataract, and a putative causal effect of type 2 diabetes on cataract particularly in East Asians. Lower expression of two novel candidate genes *MIR4453HG* and *KCNK17* were identified to be possibly involved in the causality between type 2 diabetes and cataract. Our results provided theoretical fundament at the genetic level for improving early diagnosis, prevention and treatment of cataract in type 2 diabetes patients in clinical practice.

## Data availability statement

Summary statistics are publicly available at http://jenger.riken.jp/en/.

## Supplementary Data

Supplementary data are available at IJE online.

## Author contributions

YH.Y and H.Zhao designed the study. H.Zhang and X.X conducted analyses, with assistance from YH.Y, H.Zhao and A.X. H.Zhang, YH.Y, and H.Zhao wrote the manuscript. YD.Y, YH.Y, and H.Zhao supervised the study. All authors contributed to the final revision of the paper.

## Acknowledgments

The authors thank the BBJ project for making data available.

## References

1. Charvat H, Goto A, Goto M, et al. Impact of population aging on trends in diabetes prevalence: A meta[regression analysis of 160,000 Japanese adults. Journal of Diabetes Investigation 2015; 6: 533–42

2. Tan AG, Kifley A, Tham YC, et al. Six-Year Incidence of and Risk Factors for Cataract Surgery in a Multi-ethnic Asian Population: The Singapore Epidemiology of Eye Diseases Study. Ophthalmology 2018; 125: 1844–53

3. Foster PJ, Wong TY, Machin D, Johnson GJ, Seah SKL. Risk factors for nuclear, cortical and posterior subcapsular cataracts in the Chinese population of Singapore: the Tanjong Pagar Survey. Br J Ophthalmol 2003; 87: 1112–20

4. Lee SC, Wang Y, Ko GTC, Ma RCW, Chan JCN. Risk factors for cataract in Chinese patients with type 2 diabetes: evidence for the influence of the aldose reductase gene. Clinical Genetics 2010; 59: 356–9

5. Lin HJ, Huang YC, Lin JM, et al. Novel susceptibility genes associated with diabetic cataract in a Taiwanese population. Ophthalmic genetics 2013; 34: 35–42

6. Chylack LT, Jr., Wolfe JK, Singer DM, et al. The Lens Opacities Classification System III. The Longitudinal Study of Cataract Study Group. Arch Ophthalmol 1993; 111: 831–6

7. Boomsma D, Busjahn A, Peltonen L. Classical twin studies and beyond. Nature reviews Genetics 2002; 3: 872–82

8. Cartwright N. What are randomised controlled trials good for? Philosophical Studies 2009; 147: 59.

9. Pingault J-B, O’Reilly PF, Schoeler T, Ploubidis GB, Rijsdijk F, Dudbridge F. Using genetic data to strengthen causal inference in observational research. Nature Reviews Genetics 2018; 19: 566–80

10. Bulik-Sullivan BK, Loh P-R, Finucane HK, et al. LD Score regression distinguishes confounding from polygenicity in genome-wide association studies. Nature Genetics 2015; 47: 291–5

11. Shi H, Mancuso N, Spendlove S, Pasaniuc B. Local Genetic Correlation Gives Insights into the Shared Genetic Architecture of Complex Traits. The American Journal of Human Genetics 2017; 101: 737–51

12. Davies NM, Holmes MV, Davey Smith G. Reading Mendelian randomisation studies: a guide, glossary, and checklist for clinicians. Bmj 2018; 362: k601.

13. Zhu Z, Zhang F, Hu H, et al. Integration of summary data from GWAS and eQTL studies predicts complex trait gene targets. Nat Genet 2016; 48: 481–7

14. Nagai A, Hirata M, Kamatani Y, et al. Overview of the BioBank Japan Project: Study design and profile. J Epidemiol 2017; 27: S2–S8.

15. Suzuki K, Akiyama M, Ishigaki K, et al. Identification of 28 new susceptibility loci for type 2 diabetes in the Japanese population. Nat Genet 2019; 51: 379–86

16. Ishigaki K, Akiyama M, Kanai M, Takahashi A, Kamatani Y. Large-scale genome-wide association study in a Japanese population identifies novel susceptibility loci across different diseases. Nature Genetics 2020.

17. Willer CJ, Li Y, Abecasis GR. METAL: fast and efficient meta-analysis of genomewide association scans. Bioinformatics 2010; 26: 2190–1

18. Zhou W, Nielsen JB, Fritsche LG, et al. Efficiently controlling for case-control imbalance and sample relatedness in large-scale genetic association studies. Nature genetics 2018; 50: 1335–41

19. Bulik-Sullivan B, Finucane HK, Anttila V, et al. An atlas of genetic correlations across human diseases and traits. Nature Genetics 2015; 47: 1236–41

20. International HapMap C, Altshuler DM, Gibbs RA, et al. Integrating common and rare genetic variation in diverse human populations. Nature 2010; 467: 52–8

21. da Silva JS, Wowk PF, Poerner F, Santos PS, Bicalho Mda G. Absence of strong linkage disequilibrium between odorant receptor alleles and the major histocompatibility complex. Human immunology 2013; 74: 1619–23

22. Yamada M, Hiratsuka Y, Roberts CB, et al. Prevalence of visual impairment in the adult Japanese population by cause and severity and future projections. Ophthalmic Epidemiol 2010; 17: 50–7

23. Berisa T, Pickrell JK. Approximately independent linkage disequilibrium blocks in human populations. Bioinformatics 2016; 32: 283–5

24. Burgess S, Dudbridge F, Thompson SG. Combining information on multiple instrumental variables in Mendelian randomization: comparison of allele score and summarized data methods. Stat Med 2016; 35: 1880–906

25. Bowden J, Davey Smith G, Burgess S. Mendelian randomization with invalid instruments: effect estimation and bias detection through Egger regression. International journal of epidemiology 2015; 44: 512–25

26. Zhu Z, Zheng Z, Zhang F, et al. Causal associations between risk factors and common diseases inferred from GWAS summary data. Nature Communications 2018; 9: 224.

27. Bowden J, Davey Smith G, Haycock PC, Burgess S. Consistent Estimation in Mendelian Randomization with Some Invalid Instruments Using a Weighted Median Estimator. Genet Epidemiol 2016; 40: 304–14

28. Hartwig FP, Davey Smith G, Bowden J. Robust inference in summary data Mendelian randomization via the zero modal pleiotropy assumption. International journal of epidemiology 2017; 46: 1985–98

29. Morrison J, Knoblauch N, Marcus JH, Stephens M, He X. Mendelian randomization accounting for correlated and uncorrelated pleiotropic effects using genome-wide summary statistics. Nature Genetics 2020; 52: 740–7

30. O’Connor LJ, Price AL. Distinguishing genetic correlation from causation across 52 diseases and complex traits. Nature Genetics 2018; 50: 1728–34

31. Burgess S, Davey Smith G, Davies NM, et al. Guidelines for performing Mendelian randomization investigations. Wellcome Open Res 2020; 4: 186.

32. Purcell S, Neale B, Todd-Brown K, et al. PLINK: a tool set for whole-genome association and population-based linkage analyses. Am J Hum Genet 2007; 81: 559–75

33. Auton A, Abecasis GR, Altshuler DM, et al. A global reference for human genetic variation. Nature 2015; 526: 68–74

34. Byrne EM, Zhu Z, Qi T, et al. Conditional GWAS analysis to identify disorder-specific SNPs for psychiatric disorders. Molecular Psychiatry 2020.

35. Võsa U, Claringbould A, Westra H-J, et al. Unraveling the polygenic architecture of complex traits using blood eQTL metaanalysis. 2018: 447367.

36. Willer CJ, Li Y, Abecasis GR. METAL: fast and efficient meta-analysis of genomewide association scans. Bioinformatics 2010; 26: 2190–1

37. Chang C, Zhang K, Veluchamy A, et al. A Genome-Wide Association Study Provides New Evidence That CACNA1C Gene is Associated With Diabetic Cataract. Investigative ophthalmology & visual science 2016; 57: 2246–50

38. Lin HJ, Huang YC, Lin JM, Wu JY, Tsai FJ. Single-nucleotide polymorphisms in chromosome 3p14.1- 3p14.2 are associated with susceptibility of Type 2 diabetes with cataract. Molecular Vision 2010; 16: 1206–14

39. Landrum MJ, Lee JM, Benson M, et al. ClinVar: improving access to variant interpretations and supporting evidence. Nucleic Acids Res 2018; 46: D1062–d7.

40. Rappaport N, Nativ N, Stelzer G, et al. MalaCards: an integrated compendium for diseases and their annotation. Database: the journal of biological databases and curation 2013; 2013: bat018.

41. Huang T, Wang L, Bai M, et al. Influence of IGF2BP2, HMG20A, and HNF1B genetic polymorphisms on the susceptibility to Type 2 diabetes mellitus in Chinese Han population. Bioscience reports 2020; 40.

42. El-Khairi R, Vallier L. The role of hepatocyte nuclear factor 1*β* in disease and development. Diabetes, obesity & metabolism 2016; 18 Suppl 1: 23–32

43. Xue A, Wu Y, Zhu Z, et al. Genome-wide association analyses identify 143 risk variants and putative regulatory mechanisms for type 2 diabetes. Nat Commun 2018; 9: 2941.

44. Zhu Z, Zheng Z, Zhang F, et al. Causal associations between risk factors and common diseases inferred from GWAS summary data. Nature communications 2018; 9: 1–12

45. Ismail-Beigi F, Craven T, Banerji MA, et al. Effect of intensive treatment of hyperglycaemia on microvascular outcomes in type 2 diabetes: an analysis of the ACCORD randomised trial. Lancet 2010; 376: 419–30

46. Patel A, MacMahon S, Chalmers J, et al. Intensive blood glucose control and vascular outcomes in patients with type 2 diabetes. The New England journal of medicine 2008; 358: 2560–72

47. Shichiri M, Kishikawa H, Ohkubo Y, Wake N. Long-term results of the Kumamoto Study on optimal diabetes control in type 2 diabetic patients. Diabetes care 2000; 23 Suppl 2: B21–9

48. Sun BB, Maranville JC, Peters JE, et al. Genomic atlas of the human plasma proteome. Nature 2018; 558: 73–9

49. Klarin D, Damrauer SM, Cho K, et al. Genetics of blood lipids among ~300,000 multi-ethnic participants of the Million Veteran Program. Nat Genet 2018; 50: 1514–23

50. Gannon MC, Nuttall FQ, Saeed A, Jordan K, Hoover H. An increase in dietary protein improves the blood glucose response in persons with type 2 diabetes. The American journal of clinical nutrition 2003; 78: 734–41

51. Krauss RM. Lipids and lipoproteins in patients with type 2 diabetes. Diabetes care 2004; 27: 1496–504

52. Delcourt C, Dupuy AM, Carriere I, Lacroux A, Cristol JP. Albumin and transthyretin as risk factors for cataract: the POLA study. Archives of ophthalmology (Chicago, Ill: 1960) 2005; 123: 225–32

53. Betzler BK, Rim TH, Sabanayagam C, Cheung CMG, Cheng CY. High-Density Lipoprotein Cholesterol in Age-Related Ocular Diseases. Biomolecules 2020; 10.

54. He L, Ma Q, Wang Y, et al. Association of variants in KCNK17 gene with ischemic stroke and cerebral hemorrhage in a Chinese population. Journal of stroke and cerebrovascular diseases: the official journal of National Stroke Association 2014; 23: 2322–7

55. Martín-Timón I, Sevillano-Collantes C, Segura-Galindo A, Del Cañizo-Gómez FJ. Type 2 diabetes and cardiovascular disease: Have all risk factors the same strength? World journal of diabetes 2014; 5: 444–70

56. Nemet AY, Vinker S, Levartovsky S, Kaiserman I. Is cataract associated with cardiovascular morbidity? Eye (London, England) 2010; 24: 1352–8

