## Supplementary material for "The putative causal effect of type 2 diabetes in risk of cataract: a Mendelian randomization study in East Asian": supp_fig

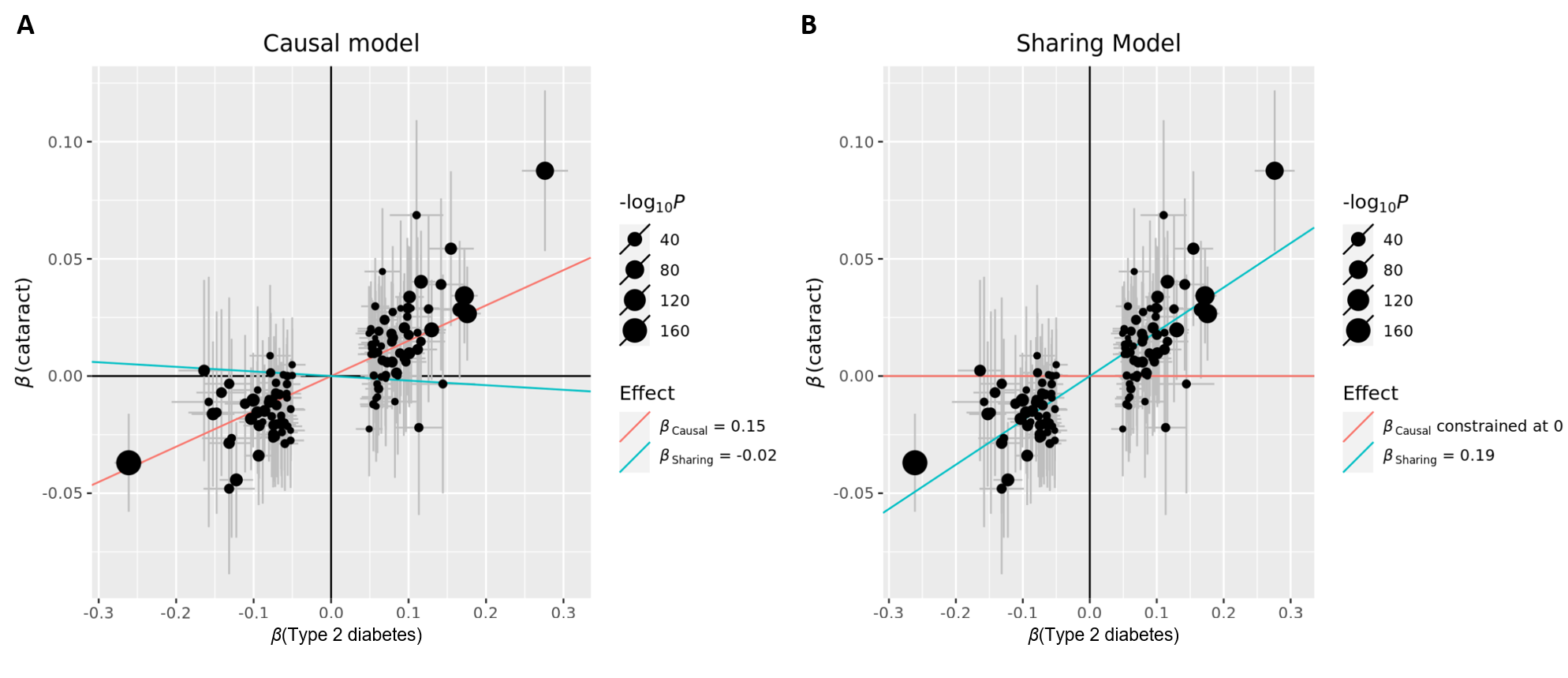

Figure S1. Comparison of CAUSE fitted causal model (left) and sharing model (right) using type 2 diabetes as exposure and cataract as outcome. Error bars stand for the 95% confidence intervals (CIs) of the SNP effect sizes based on the GWAS of type 2 diabetes (x-axis) and cataract (y-axis). Blue line represents the estimated sharing effect between type 2 diabetes and cataract. Red line represents the estimated causal effect of type 2 diabetes on cataract, which is constrained at 0 according to the sharing model. CAUSE: causal analysis using summary effect estimates.

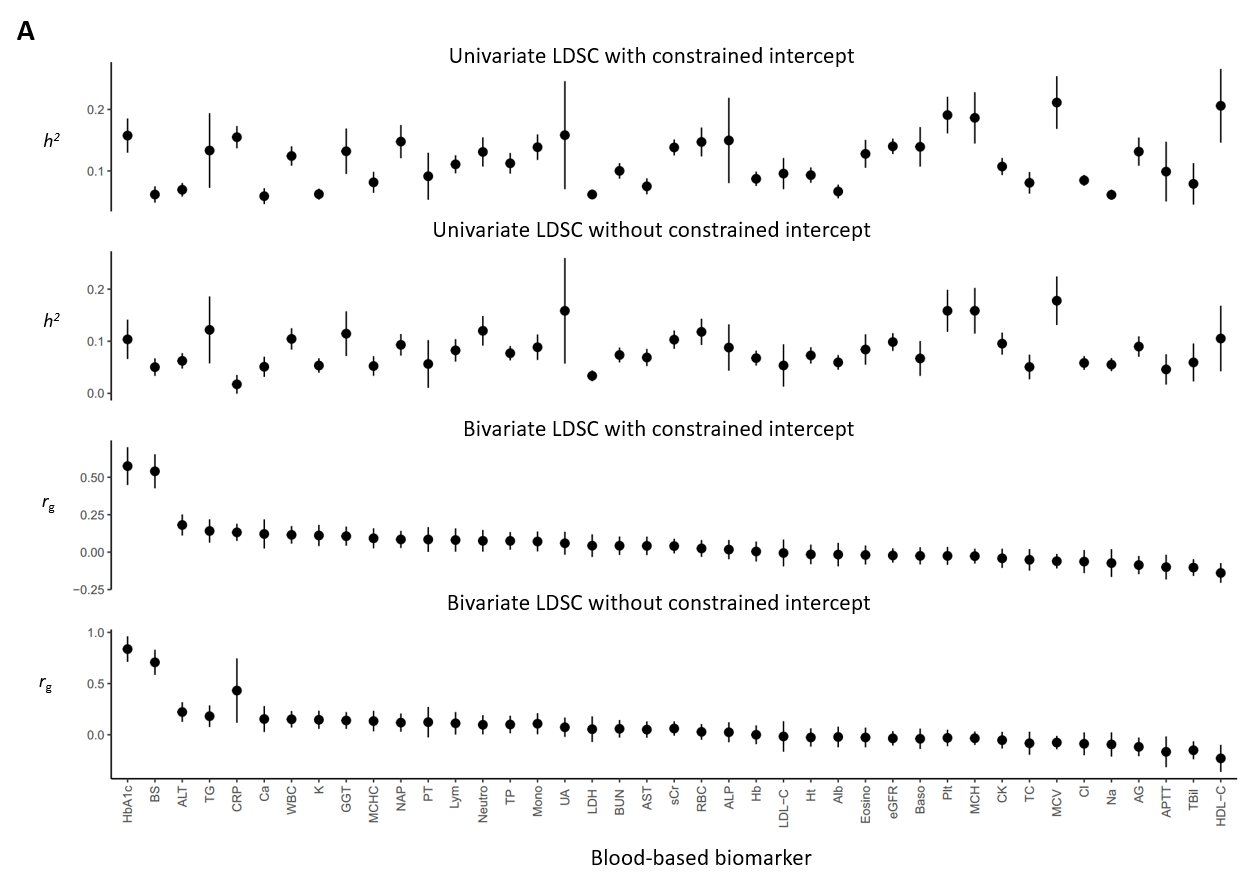

Figure S2. The estimated heritability (*h*^2^) of 41 blood-based biomarkers (summarized in Table S4) and their genetic correlation (*r*_g_) with type 2 diabetes estimated by LDSC. Error bars stand for the 95% CIs of the estimates. LDSC: linkage disequilibrium score regression.

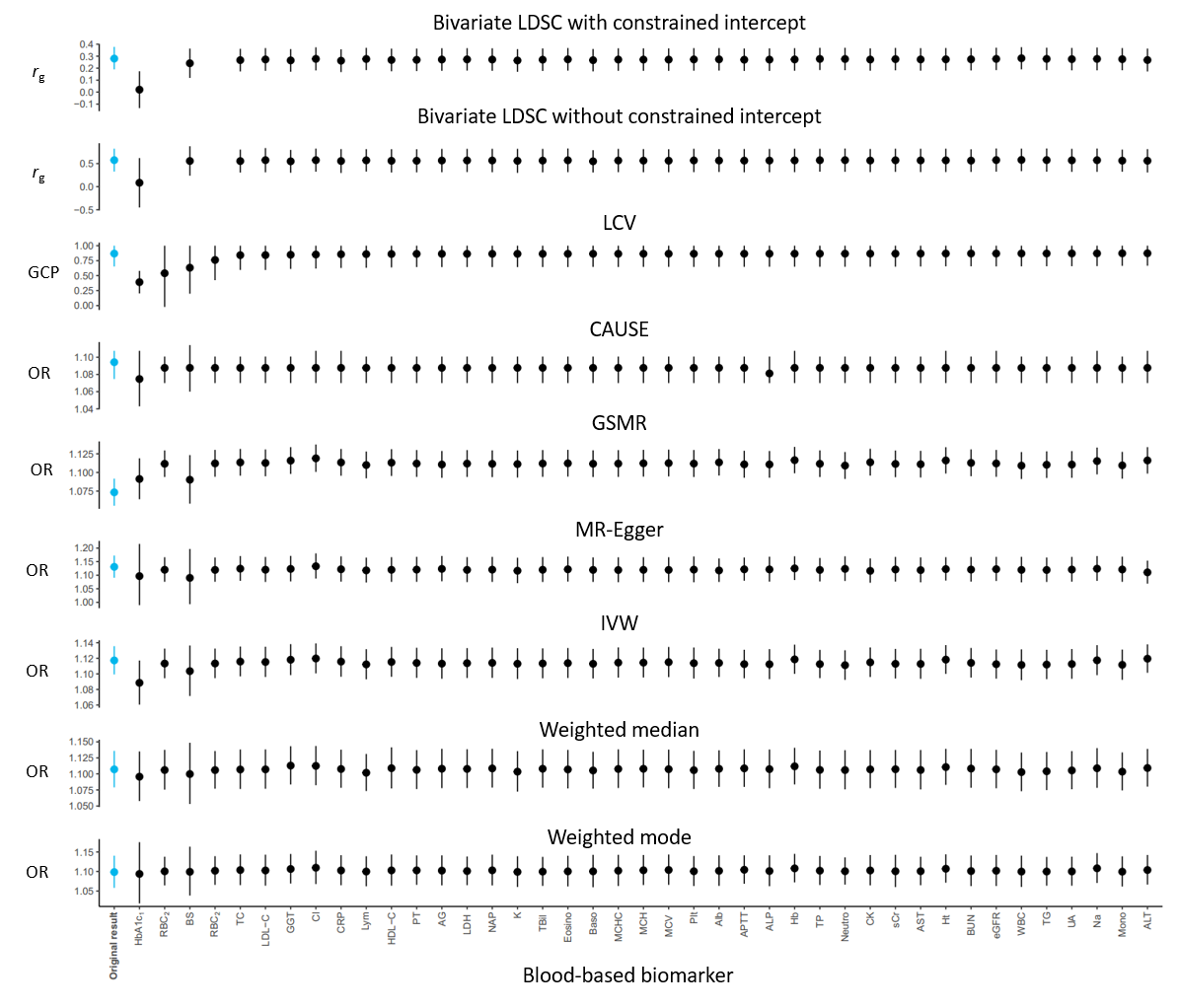

Figure S3. The genetic correlation (*r*_g_), genetic causality proportion (GCP), and liability-scale causal effect between blood-based biomarker (summarized in Table S4) adjusted type 2 diabetes (using mtCOJO, see Supplementary note for more details) and adjusted cataract estimated by LDSC and MR or MR-equivalent approaches. Error bars represent the 95% CIs of the estimates. LDSC: linkage disequilibrium score regression; LCV: latent causal variable; CAUSE: causal analysis using summary effect estimates; GSMR: generalized summary-data-based Mendelian randomization; IVW: inverse variance weighted. mtCOJO: multi-trait-based conditional and joint analysis.

^1^ Methods (except LDSC, LCV, and CAUSE) utilized ‘proxy’ independent instrumental SNPs with GWAS p-value <1×10^-5^. ^2^ No LDSC analysis performed because of the negative heritability estimates.
