## Supplementary material for "The putative causal effect of type 2 diabetes in risk of cataract: a Mendelian randomization study in East Asian": supp_note

Authors and affiliations:

Haoyang Zhang^a,b^, Xuehao Xiu^b^, Angli Xue^c^, Yuedong Yang^a,*^, Yuanhao Yang^c,d,*^, Huiying Zhao^b,*^

^a^ School of Data and Computer Science, Sun Yat-sen University, Guangzhou, China

^b^ Department of Medical Research Center, Sun Yat-sen Memorial Hospital; Guangdong Provincial Key Laboratory of Malignant Tumor Epigenetics and Gene Regulation; Guangzhou, China

^c^ Institute for Molecular Bioscience, The University of Queensland, Brisbane, QLD, Australia

^d^ Mater Research, Translational Research Institute, Brisbane, QLD, Australia

Investigations of the role of blood-based biomarkers in the causal relationship of type 2 diabetes on cataract

Data Source

We downloaded the genome-wide association studies (GWAS) summary statistics of 41 blood-based biomarkers ^1^ from the BioBank Japan Project (BBJ; <http://jenger.riken.jp/en/>), The GWAS of blood-based biomarkers were obtained by a linear model of additive allelic effects with sample size ranging from 37,767 to 143,658, adjusted by age, sex, and top ten principal components. These GWAS were based on the hg19 coordinate.

Methods

*Quality control and genotype imputation to the UK Biobank Chinese cohort*

We first generated our in-house East Asian genotype reference from the UK biobank Chinese cohort. Before imputation, we did some quality control (QC) steps to exclude the low-quality SNPs and individuals. SNPs were removed if they 1. had a missing rate >20%, 2. had minor allele frequency (MAF) <0.05, or 3. deviated from the Hardy–Weinberg equilibrium (p-value<1×10^-10^). Individuals were removed if they 1. deviated ±3 times of standard error from the sample mean of average heterozygosity rates, 2. had sex discrepancy on the heterozygosity of X chromosome, or 3. had pi‐hat (i.e., second-degree relatives) >0.2 with other individuals.

Next, the QCed genotype data were imputed according to the 1000 Genomics Phase 3 reference panel with minimac4 ^2^ using Michigan imputation server ^3^ (<https://imputationserver.sph.umich.edu/index.html>). We also conducted post-imputation QC. SNPs were removed if they 1. have imputation quality score (Rsq) < 0.3, 2. have missing rate > 20%, 3. have MAF < 0.05, 4. deviate from Hardy – Weinberg equilibrium (p-value < 1 × 10^-10^), or (5) in the histocompatibility complex (MHC) region (chromosome 6: 28 477 797–33 448 354). Individuals were removed if they (1) deviate ± 3 × SE from the sample mean of average heterozygosity rates, or (2) have pi‐hat (i.e., second-degree relatives) > 0.2 with other individuals.

*Exploration of mediated blood-based biomarkers on the causality of type 2 diabetes on cataract*

We investigated if any blood-based biomarkers might influence the causality of type 2 diabetes on cataract. The analysis was performed by the multi-trait-based conditional & joint analysis (mtCOJO) ^4^ that adjusted GWAS data of both exposure (e.g., type 2 diabetes) and outcome (e.g., cataract) for each of 41 blood-based biomarker GWAS, according to our in-house Chinese reference described above. We re-estimated the genetic correlation and potential causal relationship between the conditional GWAS of exposure and outcome using LDSC and seven MR and MR-equivalent methods, respectively. In the MR analysis, if there were <10 instrumental SNPs of exposure after conditioning on a blood-based biomarker, we selected the ‘proxy’ instrumental SNPs by releasing the GWAS p-value threshold to 1×10^-5^. If the estimates became significantly weaker or stronger compared to the initial LDSC and MR results, a blood-based biomarker was considered to be significantly up- or down-regulating the causal relationship between type 2 diabetes and cataract.

Reference

1.	Kanai M, Akiyama M, Takahashi A, et al. Genetic analysis of quantitative traits in the Japanese population links cell types to complex human diseases. *Nat Genet* 2018; **50**: 390-400.

2.	Howie B, Fuchsberger C, Stephens M, Marchini J, Abecasis GR. Fast and accurate genotype imputation in genome-wide association studies through pre-phasing. *Nat Genet* 2012; **44**: 955-9.

3.	Das S, Forer L, Schönherr S, et al. Next-generation genotype imputation service and methods. *Nat Genet* 2016; **48**: 1284-7.

4.	Zhu Z, Zheng Z, Zhang F, et al. Causal associations between risk factors and common diseases inferred from GWAS summary data. *Nature communications* 2018; **9**: 1-12.
